# Modulating the Unfolded Protein Response with ISRIB Mitigates Cisplatin Ototoxicity

**DOI:** 10.1101/2023.10.17.562797

**Authors:** Jiang Li, Stephanie L. Rouse, Ian R. Matthews, Elliott H. Sherr, Dylan K. Chan

**Author notes:** These two authors share first position (Jiang Li, Stephanie L. Rouse). These two authors share last position (Elliott H. Sherr, Dylan K. Chan). To whom correspondence should be addressed: Dylan K. Chan, MD, PhD Department of Otolaryngology-Head and Neck Surgery, UCSF 513 Parnassus Ave, Rm 719 San Francisco, CA 94143.

## Abstract

Cisplatin is a commonly used chemotherapy agent with a nearly universal side effect of sensorineural hearing loss. The cellular mechanisms underlying cisplatin ototoxicity are poorly understood. Efforts in drug development to prevent or reverse cisplatin ototoxicity have largely focused on pathways of oxidative stress and apoptosis. An effective treatment for cisplatin ototoxicity, sodium thiosulfate (STS), while beneficial when used in standard risk hepatoblastoma, is associated with reduced survival in disseminated pediatric malignancies, highlighting the need for more specific drugs without potential tumor protective effects. The unfolded protein response (UPR) and endoplasmic reticulum (ER) stress pathways have been shown to be involved in the pathogenesis of noise-induced hearing loss and cochlear synaptopathy *in vivo*, and these pathways have been implicated broadly in cisplatin cytotoxicity. This study sought to determine whether the UPR can be targeted to prevent cisplatin ototoxicity. Neonatal cochlear cultures and HEK cells were exposed to cisplatin and UPR-modulating drugs, and UPR marker gene expression and cell death measured. Treatment with ISRIB, a drug that activates eif2B and downregulates the pro-apoptotic PERK/CHOP pathway of the UPR, was tested in an *in vivo* mouse model of cisplatin ototoxicity and well as a head and neck squamous cell carcinoma (HNSCC) cell-based assay of cisplatin cytotoxicity. Cisplatin exhibited a biphasic, non-linear dose-response of cell death and apoptosis that correlated with different patterns of UPR marker gene expression in HEK cells and cochlear cultures. ISRIB treatment protected against cisplatin-induced hearing loss and hair-cell death, but did not impact cisplatin’s cytotoxic effects on HNSCC cell viability, unlike STS. These findings demonstrate that targeting the pro-apoptotic PERK/CHOP pathway with ISRIB can mitigate cisplatin ototoxicity without reducing anti-cancer cell effects, suggesting that this may be a viable strategy for drug development.

## Introduction

Cisplatin is an effective chemotherapeutic agent used to treat many malignant solid tumors, yet treatment is associated with ototoxicity. Irreversible sensorineural hearing loss affects 75-100% of patients treated with cisplatin, with young children more susceptible than adults (1–4). Despite some efficacy for sodium thiosulfate in the treatment of cisplatin ototoxicity in non-metastatic hepatoblastoma in children (5,6), significant concerns have been raised that STS reduces overall survival especially in metastatic solid tumors, possibly due to interfering with cisplatin’s anti-cancer effects (7). There remains, therefore, a significant need for more specific pharmacological intervention to prevent ototoxicity while preserving cancer-cell cytotoxicity.

The cellular mechanisms underlying cisplatin ototoxicity are poorly understood. Upon exposure, cisplatin is taken up into hair cells principally through the copper transporter Ctr1 (8) and binds DNA, leading to generation of excessive ROS, inflammation, transcriptional and cell cycle arrest, ultimately causing apoptosis and necrosis of sensory hair cells as well as cells in the stria vascularis and spiral ganglion (9–13). Efforts in drug development to prevent or reverse cisplatin ototoxicity have largely focused on pathways of oxidative stress and apoptosis (14). Exogenous administration of antioxidants mitigates hearing loss, but antioxidants are not a viable solution as antioxidant stress is also harmful to the cochlea (14).

There is extensive crosstalk between oxidative stress and endoplasmic reticulum (ER) stress (15), as up to 25% of ROS are generated in the ER (16), suggesting that ER dysfunction may be crucial to understanding cisplatin-induced cochlear cell apoptosis. ER stress occurs when improperly folded proteins accumulate in the ER lumen. In response, the unfolded protein response (UPR) is activated to either restore ER proteostasis, in the case of more transient stress, or induce apoptosis in severe stress (17).

Cisplatin induces ER stress (18); it directly interacts with proteins (19) and promotes their unfolding (20), and damage done to other organelles by cisplatin may act as an indirect inducer of ER stress (21). Cisplatin ototoxicity is correlated with cytosolic protein synthesis inhibition (22). In a rat model of cisplatin ototoxicity, BiP expression was broadly upregulated, indicating induction of ER stress. Enhancement of ER chaperone function and proteostasis with tauroursodexoycholic acid protected against cisplatin ototoxicity *in vivo*, indicating the UPR is a viable target for therapy against cisplatin ototoxicity (23). However, while ER stress has been implicated, the precise role of the UPR in cisplatin ototoxicity and cytotoxicity is not fully understood.

Our group implicated the UPR and ER stress in rapidly progressive genetic hearing loss due to absence of the novel deafness gene, TMTC4, as well as in noise induced hearing loss (NIHL) and cochlear synaptopathy (24,25). We demonstrated that the UPR is activated within the cochlea of adult mice within two hours of noise exposure, and that treating mice with ISRIB (Integrated Stress Response inhIBitor), a small molecule activator of eIF2B that results in downregulation of CHOP and the pro-apoptotic arm of the UPR (26), protects against NIHL and synapse loss.

Based on our findings implicating the UPR in genetic and noise-induced hearing loss, and the work of others implicating ER stress and the UPR in cisplatin ototoxicity (27,28), we sought to further explore the role of the UPR in the pathogenesis and treatment of cisplatin-induced ototoxicity. In this study, we extend our finding that the UPR is involved in NIHL, and that ISRIB can prevent NIHL, by showing that the UPR also mediates cisplatin ototoxicity. Most significantly, we demonstrate that ISRIB reduces cisplatin ototoxicity *in vivo* without affecting cisplatin’s cytotoxic effects on head-and-neck squamous cell carcinoma (HNSCC) cells *in vitro*. Our findings support the idea that the UPR is a common early pathway in NIHL and cisplatin ototoxicity and may be an appealing target for therapeutic intervention.

## Results

### Non-linear cisplatin-induced cell death is correlated with UPR gene expression

To explore the mechanistic relationship between cisplatin cytotoxicity and UPR gene expression, we first measured UPR marker gene expression in response to a broad range of cisplatin doses in HEK cells. UPR apoptosis/cell fate is regulated predominately through the actions of the IRE1 and PERK arms sending pro-homeostatic and pro-apoptotic signals, respectively (**Figure 1A**). Death receptor 5 (DR5) integrates the opposing UPR signals of these two arms to control ER-stress-induced apoptosis: PERK/CHOP activity induces *DR5* transcription leading to formation of caspase 8 activating complex and ultimately apoptosis, whereas IRE1α/S-XBP1 promotes *DR5* mRNA decay, giving the cell time to restore homeostasis (29). Thus, DR5 levels are a measure of the persistence of ER stress and likelihood of apoptosis and are controlled by the opposing activities of the PERK/CHOP and IRE1α/S-XBP1 pathways.

**Figure 1.**
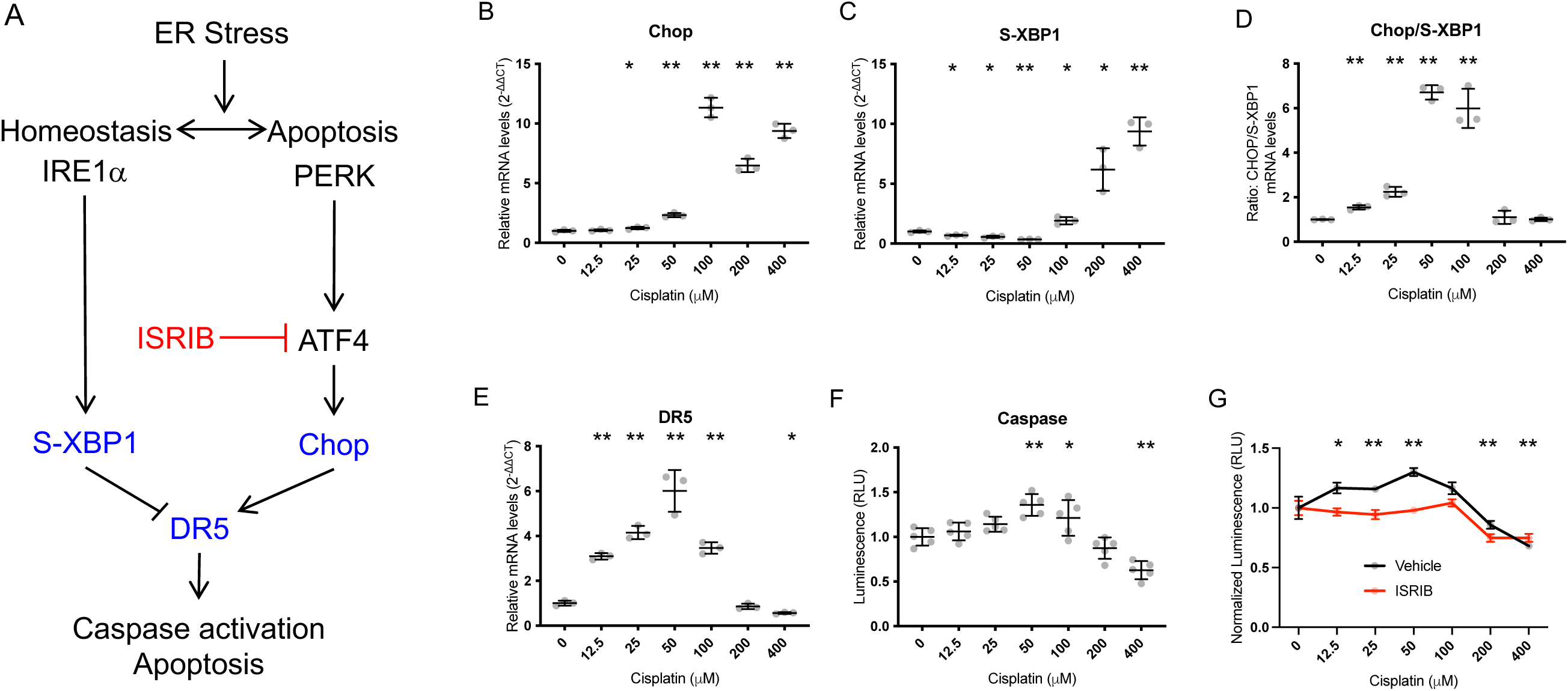
Non-linear cisplatin dose-response of UPR gene expression and apoptosis in HEK cells. **A.** In the unfolded protein response (UPR), two major pathways, labeled by their initiating effectors (IRE1a, and PERK) respond to accumulation of misfolded or unfolded proteins by facilitating homeostasis or, if the load is too great, apoptosis. The downstream effectors S-XBP1 and Chop are commonly used markers of the homeostatic (S-XBP1) and apoptotic (Chop) pathways. Depending on persistence and severity of stress, indicated by DR5, which integrates S-XBP1 and Chop signals, the UPR halts translation and allows for an adaptive response and return to proteostasis, or terminal response ending in apoptosis. ISRIB inhibits the PERK pathway by activating eiF2B and inhibiting ATF4. **B-E. UPR gene expression**. Wild-type HEK cells were treated with multiple doses of cisplatin, and mRNA levels of Chop, S-XBP1, the ratio of Chop/S-XBP1, and DR5 were measured by qPCR and normalized to GAPDH by the 2^-ΔΔCT^ method. **F. Apoptosis.** Caspase 3/7 activity was quantified using Caspase-Glo, demonstrating a non-linear cisplatin dose-dependence of HEK cells that matches DR5 expression. **B-F.** Data are means ± SEM, with individual data shown as grey dots. * p < 0.05; ** p < 0.001 by one-way ANOVA followed by pairwise t-test relative to control (0 μM cisplatin). Each graph is from a single experiment with 3 (**B-E**) or 5 (**F**) independent samples for each cisplatin dose. The experiment was repeated three times with identical results. **G. ISRIB treatment in cisplatin-treated cells reduces apoptosis.** HEK cells treated for 40 h with cisplatin were additionally treated with 3 doses (at 0, 16, and 24 h) of 0.2 μM ISRIB (red), or vehicle (black), and Caspase 3/7 activity measured. Increasing cisplatin concentration induced an initial increase, then decrease, in apoptosis in the absence of ISRIB, consistent with the non-linear pattern of hair-cell death seen in cisplatin-treated cochlear cultures. At low cisplatin doses, ISRIB attenuated cisplatin-induced apoptosis and eliminated cisplatin dose-dependence; at high cisplatin doses, ISRIB had no effect. Data are means ± SEM, with individual values as pink or grey dots. *p<0.05; **p<0.01 compared to no drug. N=3 for each condition.

Surprisingly, HEK cells exhibited a biphasic, bell-shaped response of UPR markers Chop, S-XBP1, and DR5 with increasing cisplatin dose, with a peak in expression of pro-apoptotic Chop and trough in expression of pro-homeostatic S-XBP1 at 50-100 µM cisplatin (**Figure 1B-C**). The Chop/S-XBP1 ratio and DR5 expression were similarly elevated in this range, consistent with integration of the Chop and S-XBP1 pathways towards apoptosis (**Figure 1D-E**). In accordance with this, we observed a peak in apoptosis, measured by caspase 3/7 activity, at 50 µM cisplatin (**Figure 1F**).

Observation of this correlation of UPR gene expression and apoptosis across this non-linear, bi-phasic dose response led us to hypothesize that the UPR is causally involved in, and may be targeted to reduce, cisplatin-induced apoptosis. To test this, we treated cells with 0.2 μM ISRIB, an eIF2B activator that inhibits the PERK/CHOP arm of the UPR (26) (**Figure 1A**) and measured the effect on apoptosis across the same range of cisplatin doses. ISRIB reduced cisplatin-induced apoptosis at low cisplatin doses but had no effect at high cisplatin doses (**Figure 1G**), suggesting that it may be an effective drug to reduce cisplatin ototoxicity.

### Cisplatin treatment induces the Unfolded Protein Response in the cochlea *in vitro*

ER stress and the UPR has been broadly implicated in cytotoxicity induced by cisplatin treatment (18). We treated neonatal cochlear explant cultures with 100 µM cisplatin for 20h and performed RNAScope to detect Chop. Cisplatin-induced Chop expression was detected principally in cells of the stria vascularis and inner and outer sulcus supporting cells, but not in the hair-cell region (**Figure 2**).

**Figure 2.**
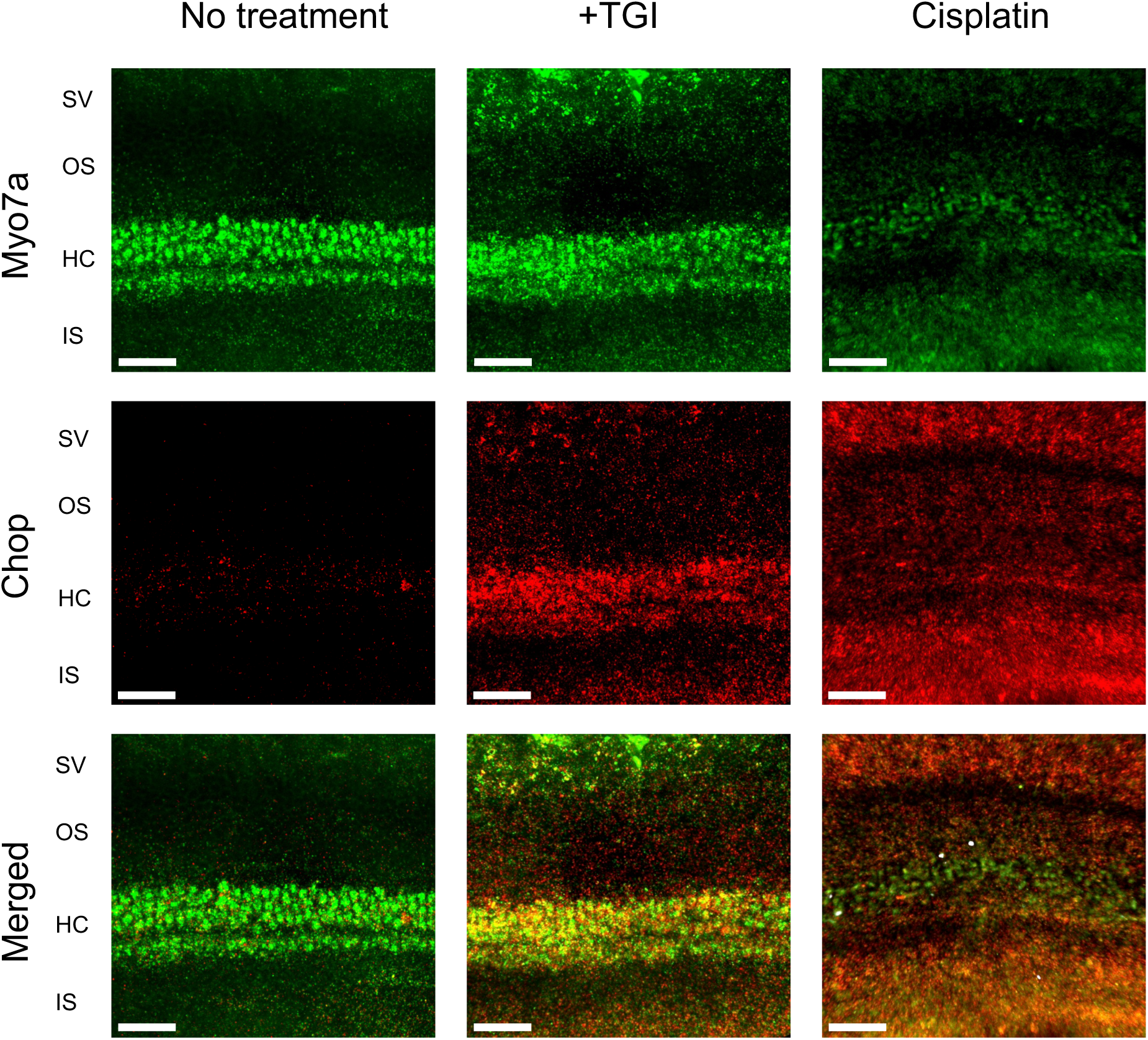
Chop expression detected in cochlear cultures in response to TG and cisplatin. RNAscope probing for Myo7a (green) and Chop (red) in P2 C57 cochlear cultures after 2 h treatment with thapsigargin (TG, a known inducer of the UPR, middle) shows increased expression of Chop in TG-treated cultures, particularly in the hair-cell region (HC). Cultures treated with 100 μM cisplatin (right) for 20h demonstrated upregulation of Chop in stria vascularis (SV) and supporting cells of the inner and outer sulcus (IS/OS), but not hair cells. Scale bars: 50 μm.

We then sought to quantify the effect of cisplatin on UPR markers in the cochlea. Explant cultures of the P3-5 neonatal cochleae from wild-type C57BL/6J mice were treated with escalating concentrations of cisplatin for 20h and then subjected to whole-mount immnunohistochemistry against Myo7a, a hair-cell marker. Loss of inner and outer hair cells occurred with increasing cisplatin dose up to 100 µM; with higher doses, hair cells were paradoxically preserved up to 1000 µM cisplatin (**Figure 3A**).

**Figure 3.**
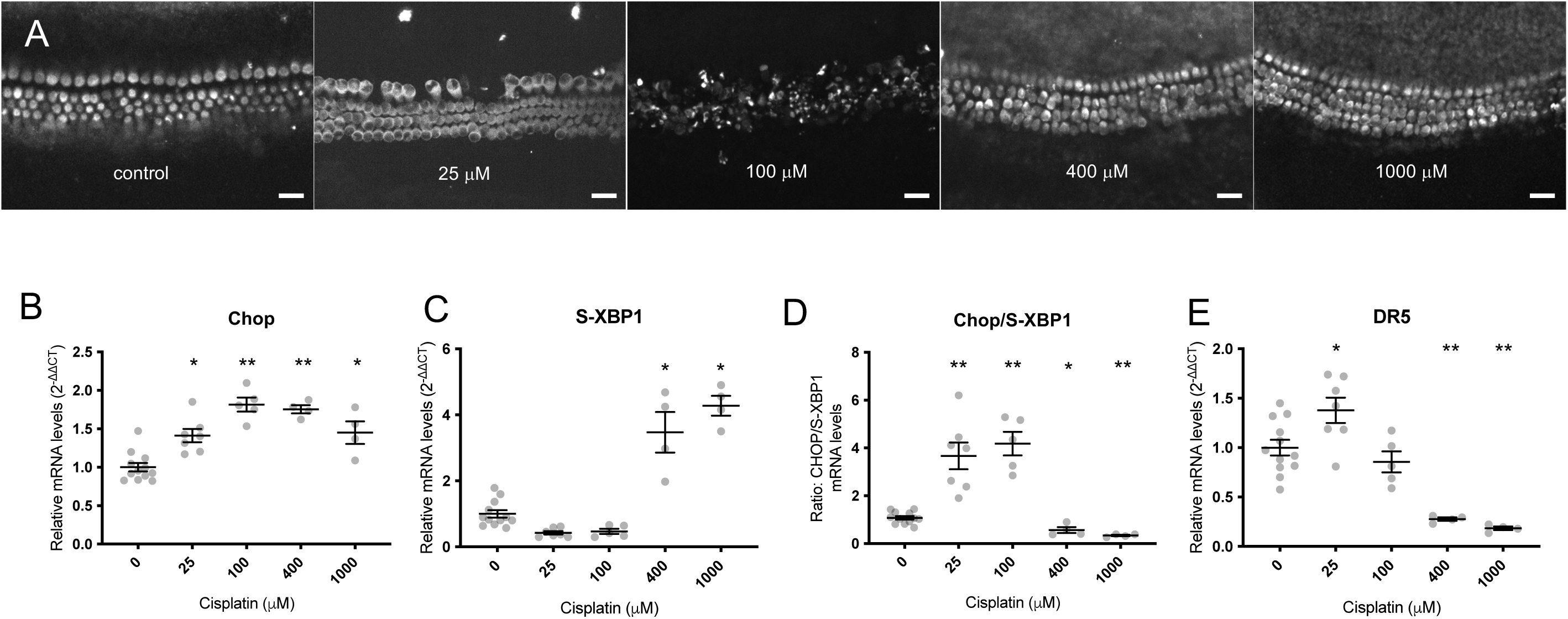
UPR modulation in cisplatin-treated cochleae. **A.** Non-linear hair-cell death in response to cisplatin. Bell-shaped dose response in hair-cell death is observed in P3 WT C57BL/6J organotypic cochlear cultures treated with cisplatin at increasing doses (0-1000 μM) for 24 h then fixed and stained with anti-Myo7a to detect hair cells. At low concentrations of cisplatin, hair-cell death increases with increasing cisplatin concentration, but above 100 μM, cisplatin-treated cochleae had preserved hair cells. **B-E.** Non-linear UPR gene expression in cochlear cultures quantified by qPCR after 6 h exposure to cisplatin. Chop (**C**), a marker of the pro-apoptotic arm of the UPR, and S-XBP1 (**C**), a pro-homeostatic UPR marker, displayed inverse non-linear dose-dependences. **D.** The ratio of Chop/S-XBP1 demonstrated elevated balance towards pro-apoptotic Chop expression consistent with the pattern of hair-cell death seen in **A.** The dose-dependency of this ratio was consistent with the expression pattern of DR5, a downstream marker that integrates the Chop and S-XBP1 signals and signals towards apoptosis. **B-E**. Data are 2^-ΔΔCT^ measurements compared internally to GAPDH and normalized to control, non-treated cultures from the same experimental run, and are presented as means ± SEM, with individual data shown as grey dots. *p < 0.05; **p < 0.001 by one-way ANOVA relative to control (0 μM cisplatin). Scale bars: 20 μm.

Multiple groups have observed this bell-shaped dose-response of hair-cell death upon exposure of neonatal cochlear cultures to cisplatin: low cisplatin concentrations induce hair-cell death that increases with increasing cisplatin dose, whereas high cisplatin concentrations cause minimal hair-cell loss (22, 27–28). Based on our findings in HEK cells (**Figure 1**), we hypothesized that the UPR may play a role in this response.

We performed qPCR on murine cochlear explant cultures exposed to the same escalating doses of cisplatin for 6h, and measured Chop, S-XBP1, and DR5 expression. Expression of these three markers exhibited a similar bell-shaped dose-response curve with inflection at cisplatin doses of 25-100 µM cisplatin dose (**Figure 3B-E**). The highest expression level of the pro-apoptotic marker Chop was seen at a cisplatin dose of 100 µM, whereas the lowest expression level of the homeostatic marker S-XBP1 was seen at cisplatin doses of 25 and 100 µM. The ratio of Chop to S-XBP1, which reflects the balance between pro-apoptotic and pro-homeostatic arms of the UPR, demonstrated the same bell-shaped dose-response, and DR5, which integrates the two pathways and therefore reflects this Chop/S-XBP1 ratio, was maximally expressed at 25 µM cisplatin. The similarity between the unusual bell-shaped dose-response relationships of cisplatin with hair-cell death and pro-apoptotic UPR marker expression suggests that UPR activity may be involved in mediating cisplatin-induced hair-cell death.

### ISRIB protects against cisplatin-induced hearing loss and hair cell loss *in vivo*

These *in vitro* experiments suggest that cisplatin may induce hair-cell death via ER stress and induction of the UPR, and that ISRIB, an eIF2B activator, is effective at reducing cisplatin-induced apoptosis. Based on previous work suggesting that targeting ER stress can reduce cisplatin ototoxicity (23) and our finding that ISRIB protects against ER stress and NIHL (24), we hypothesized that ISRIB would also be effective at mitigating cisplatin ototoxicity. To test this, we used a model of cisplatin ototoxicity in Balb/cJ mice, a strain that has been shown to have low mortality with cisplatin treatment (30). A single intraperitoneal (IP) dose of 5.5 mg/kg cisplatin administered to 8-week-old female wild-type Balb/cJ mice caused reproducible, isolated 32-kHz ABR threshold elevation at post-injection-day (PID) 21, with minimal morbidity and no mortality (**Figure 4**). Cisplatin-treated mice experienced a transient loss of 10% body weight at PID7, with full recovery to baseline by PID21, with no mortality observed (N=23), suggesting no overall impact on Balb/cJ morbidity.

**Figure 4.**
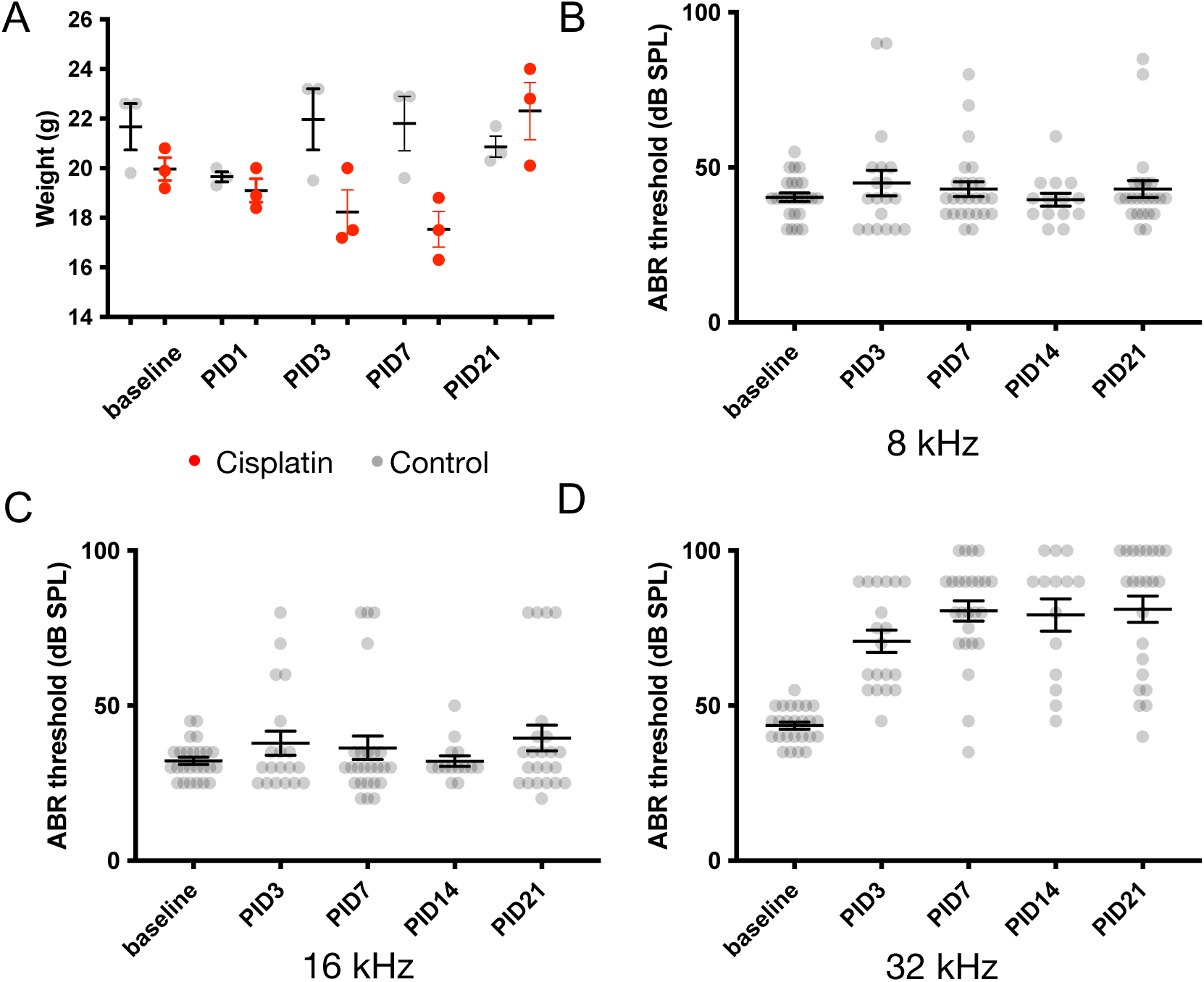
Balb/cJ model of murine cisplatin ototoxicity. A single intraperitoneal dose of 5.5 mg/kg cisplatin (CP) or saline was administered to 8-week-old female Balb/cJ mice. **A.** Compared to control mice (gray, N=3), weight decreased slightly in the CP group (red, N=3) but returned to baseline by post-injection-day (PID)21. No mortality occurred. **B-D.** ABR thresholds to 8, 16, and 32 kHz pure-tone pips were recorded at PID3, 7, 14, and 21. Data shown are means ± SEM, with individual data shown in gray dots.

10 mg/kg IP ISRIB was administered concurrent with cisplatin and serial ABR performed at PID7, 14, and 21. Cisplatin-induced 32 kHz ABR threshold elevation was reduced in ISRIB-treated mice compared with vehicle-treated controls at all 3 timepoints (**Figure 5A**, p < 0.01). 21 days after cisplatin injection (PID21), ISRIB-treated animals had a mean 32-kHz ABR threshold of 58.3 ± 21.8 (mean ± SD, N=6), compared with 89.2 ± 17.4 dB SPL for vehicle-treated controls, a statistically significant difference (p = 0.02, unpaired two-tailed t test). In fact, 32-kHz ABR ISRIB-treated animals exhibited no significant elevation in 32-kHz ABR thresholds (44.2 ± 10.7 dB SPL at baseline compared with 58.3 ± 21.8 dB SPL at PID21, p = 0.18). Overall, ISRIB treatment was associated with a 69% reduction in cisplatin-induced threshold shift compared to vehicle.

**Figure 5.**
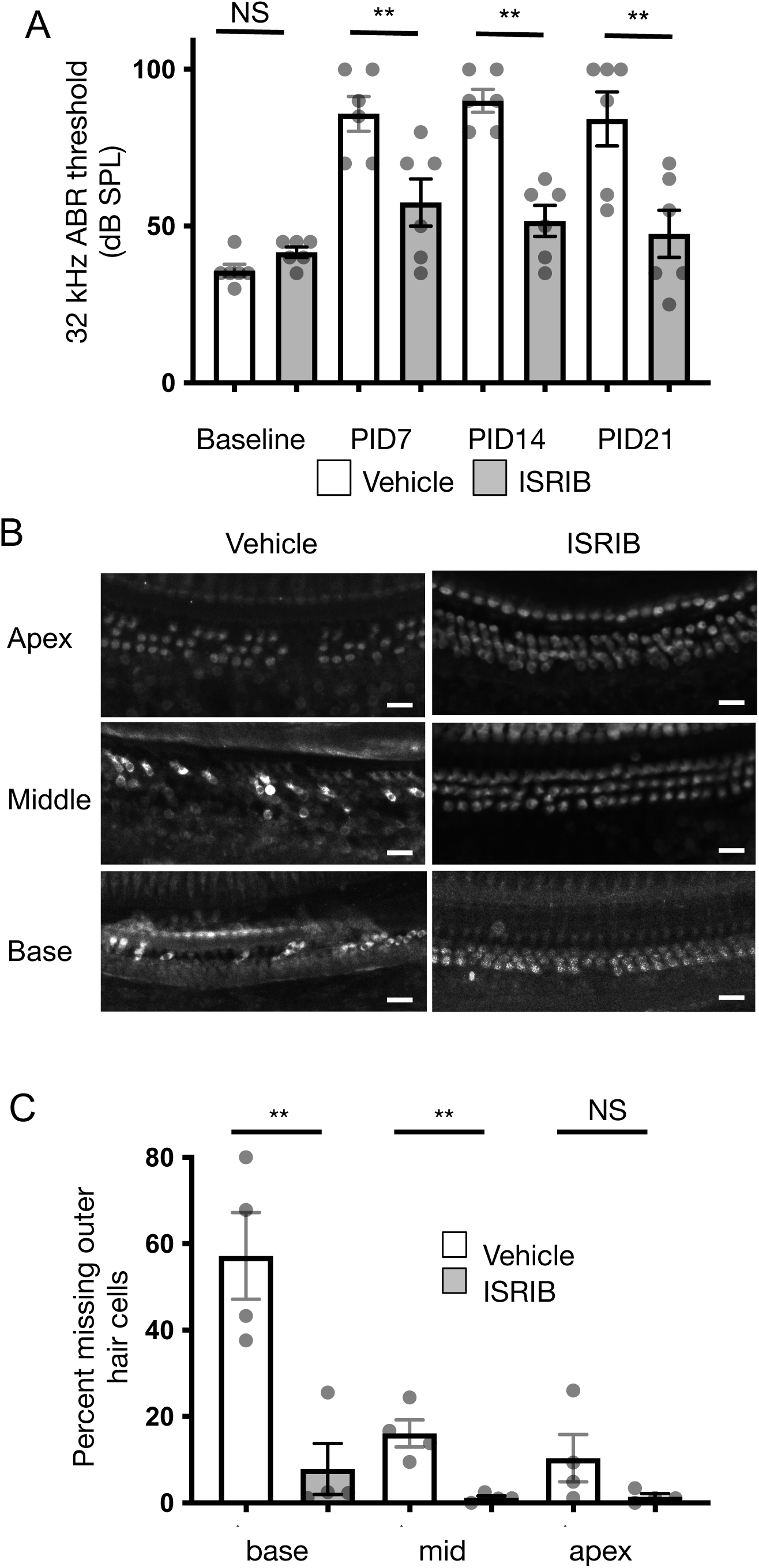
ISRIB protects against cisplatin-induced hearing and hair-cell loss. 8-week-old Balb/cJ mice were treated with a single intraperitoneal dose of 5.5 mg/kg cisplatin, and either 10 mg/kg ISRIB or vehicle. ABR thresholds to pure-tone pips were measured 7, 14, and 21 days after cisplatin treatment. After 21 days, mice were euthanized and temporal bones harvested. **A.** 32-kHz ABR threshold measurements demonstrated significant protection against cisplatin-induced hearing loss by in ISRIB-treated animals (gray) compared to controls (white). **B.** Cochlear whole mounts from the apical, middle, and basal turns were stained with Myo7a and visualized by confocal microscopy, demonstrating significant hair-cell loss in cisplatin + vehicle treated animals (left), worse in the middle and basal turns. Cochleae from animals treated with ISRIB had better hair-cell preservation after cisplatin exposure. **C.** Hair cells were quantified as the percentage of missing hair cells in a fixed-length segment from the cochlear basal, middle, or apical turns from mice 21 days after treatment with cisplatin and either ISRIB (gray, N=4) or vehicle (white, N=4). Data shown are means ± SEM, with individual data shown in grey dots. ** *p* < 0.01; NS; not significant. Scale bars: 20 μm.

To evaluate further for ototoxicity, whole-mount immunohistochemistry was performed after 21 days of cisplatin exposure to quantify outer-hair-cell (OHC) loss. Whereas 57% ± 10% (mean ± s.e.m., N=4) of basal-turn OHCs were lost in mice treated with cisplatin and vehicle, mice co-treated with ISRIB and cisplatin exhibited little to no OHC loss (7.9% ± 5.9%, N=4, p <0.01 compared to vehicle; **Figure 5B,C**). These results demonstrate that ISRIB co-treatment protects against cisplatin-induced hearing loss and OHC death in this *in vivo* model of cisplatin ototoxicity.

### ISRIB does not affect the cytotoxicity of cisplatin against cancer cells *in vitro*

A principal concern for any drug treatment for ototoxicity is potential interference with the anti-tumor effects of cisplatin. To test this, we used the FaDu head and neck squamous-cell carcinoma (HNSCC) cell line (gift of D.J.) (31). Cisplatin treatment reduces FaDu cell viability, with an IC_50_ of 25.6 μM (R^2^=0.98). Co-treatment with 20 μM z-VAD-FMK, a caspase inhibitor known to inhibit cisplatin-induced hair-cell apoptosis (32) as well as cisplatin chemotoxicity in cancer cells (33), significantly increased the IC_50_ of cisplatin (33.3 μM (R^2^=0.98); IC_50_ shift ratio re: control = 1.32 (95% CI: 1.19-1.48), **Figure 6**). Similarly, co-treatment with 3 mM STS, the only effective drug for cisplatin ototoxicity humans that also has been shown to be associated with worse survival in disseminated hepatoblastoma (7), significantly increased the IC_50_ of cisplatin (151.6 μM (R^2^=0.97); IC_50_ shift = 10.55 (9.17-12.21)). In contrast, co-treatment with 0.2 μM ISRIB, which prevented cisplatin-induced apoptosis in HEK cells (**Figure 1A**), did not affect the IC_50_ of cisplatin (27.7 μM (R^2^=0.98); IC_50_ shift = 1.01 (95% CI: 0.90-1.14)) in FaDu cells. These findings demonstrate that ISRIB does not reduce the chemo-efficacy of cisplatin in this HNSCC cell line *in vitro*.

**Figure 6.**
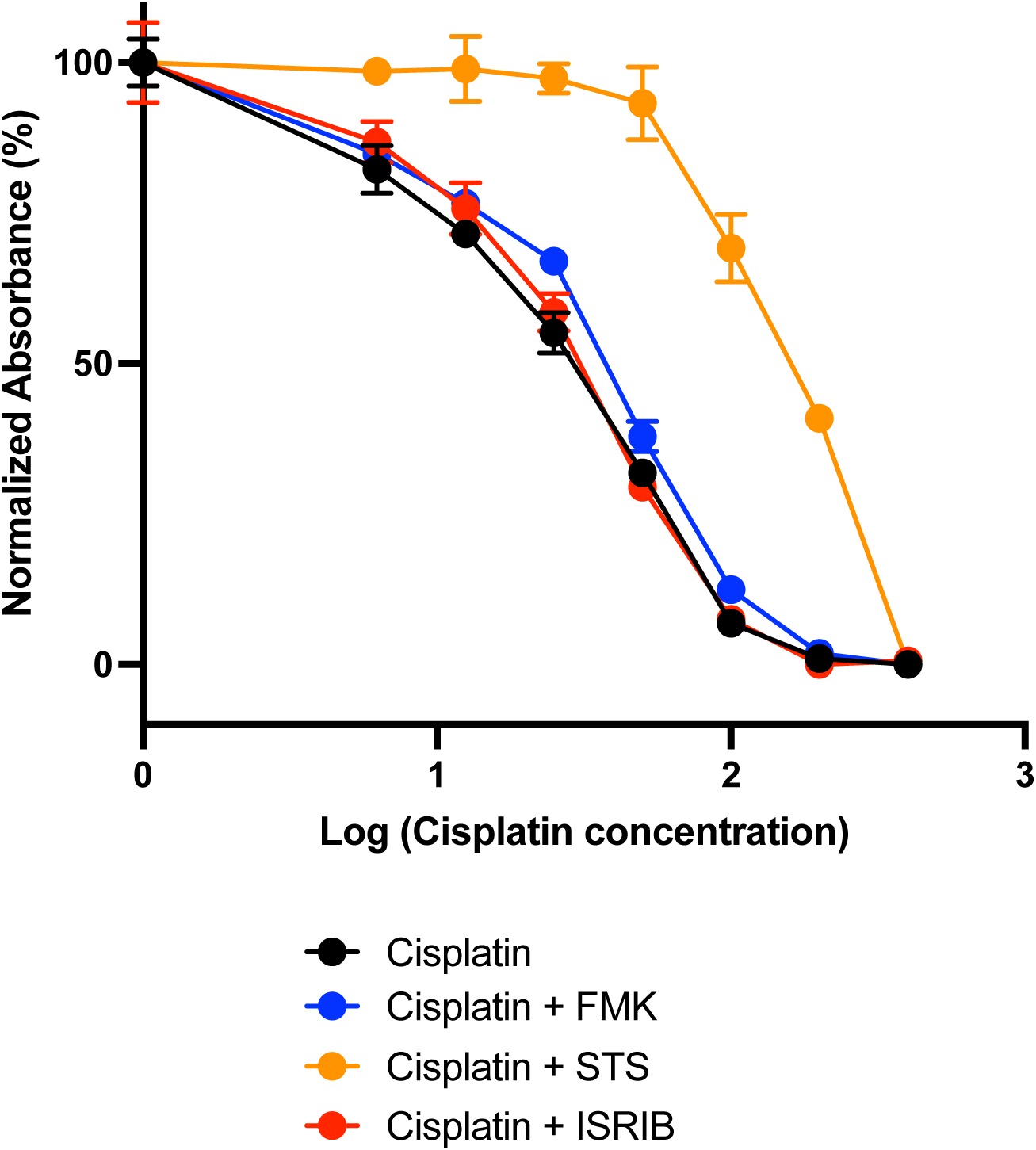
ISRIB does not interfere with cisplatin cytotoxicity against cancer cells. Viability of FaDu cells, a head and neck squamous-cell carcinoma (HNSCC) cell line, was measured in presence of escalating doses of cisplatin alone (black) or in combination with 20 μM z-VAD-FMK (blue), an apoptosis inhibitor, 3 mM STS (orange), or 0.2 μM ISRIB (red). Data shown are means ± SEM (N = 3 for each condition).

## Discussion

Cisplatin is a frequently used chemotherapeutic agent with the potentially dose-limiting and life altering side effect of sensorineural hearing loss. However, the cellular mechanism underlying cisplatin ototoxicity is poorly understood. Here, we present evidence suggesting that the UPR may play an important role in mediating cisplatin-induced ototoxicity. Using a novel mouse model of cisplatin ototoxicity that produces consistent high-frequency hearing threshold elevation with minimal morbidity, we additionally show that modulation of the UPR with ISRIB protects against cisplatin-induced hearing loss and hair-cell death. Unlike the pan-caspase inhibitor z-VAD-fmk and the cisplatin ototoxicity drug STS, ISRIB does not impair the cytotoxicity of cisplatin in a HNSCC cell line. These findings suggest that targeting eIF2b using ISRIB may be a promising strategy to treat cisplatin ototoxicity without interfering with cisplatin chemo-efficacy.

### Non-linear dose-response of cisplatin and hair-cell death

We observed a bell-shaped dose response of hair-cell death upon exposure of neonatal cochlear cultures to cisplatin: low cisplatin concentrations induce hair-cell death that increases with increasing cisplatin dose, whereas high cisplatin concentrations give rise to little or no hair-cell loss. This phenomenon has been observed previously (27–28), but the mechanism is unknown. One possibility is that cisplatin uptake may be altered at high cisplatin concentrations (27). Other studies have suggested that this phenomenon is an artifact of retention of hair cell “corpses” that retain hair-cell markers (34–35). However, a more recent study showed this is not the case: 500 μm cisplatin treatment caused arrest of protein synthesis, but subsequent washout permitted recovery of protein synthesis and hair-cell survival (22).

We observed in HEK cells and cochlear explant cultures that expression of Chop, S-XBP1, and DR5, markers of UPR activation specifically in the apoptotic, homeostatic, and integrated pathways, respectively, all displayed similar, unusual bell-shaped dose-response curves. This correlation suggests that the UPR plays a mechanistic role in cisplatin ototoxicity. The increase in S-XBP1 expression at high cisplatin concentrations, in particular, suggests that this protective response may underlie the reduction in hair-cell loss at these high concentrations. ISRIB, an eIF2B activator that suppresses the pro-apoptotic activity of the UPR, protected against cisplatin-induced apoptosis in the lower, physiologically relevant range of cisplatin doses, further supporting a mechanistic role of the UPR in cisplatin ototoxicity.

Identification of different control pathways for cisplatin-induced cytotoxicity in different cell types could be of significant value in development of drugs to protect against cisplatin ototoxicity while retaining its cytotoxic properties for cancer chemotherapy. Studies in multiple cancer cell types have suggested that the homeostatic arm of the UPR plays a central role in chemoresistance (36–38). *In vitro*, cisplatin affects cancer cells and hair cells differently: whereby cochlear hair cells exhibit the non-linear, bell-shaped dose response, cancer cell lines such as head and neck squamous cell carcinoma exhibit a typical monotonic dose-response (39). It is possible that leveraging different dose-response characteristics relating to the UPR could lead to the development of UPR-targeting drugs that protect the cochlear while retaining anti-tumor activity. Indeed, we found that ISRIB, which specifically targets eIF2B, did not alter cisplatin’s cytotoxic effect on a HNSCC cell line, providing strong initial evidence supporting this strategy. Multiple additional small molecule drugs have been developed to target different points in the homeostatic and apoptotic arms of the UPR. These can be used *in vitro* and *in vivo* to further test the hypothesis that the UPR is a control mechanism and target for treatment of cisplatin ototoxicity.

### ISRIB protects against cisplatin ototoxicity

The UPR is a potential target pathway for treatment of cisplatin ototoxicity. For this study, we refined a single-dose mouse model of acute cisplatin ototoxicity. Murine models of cisplatin ototoxicity are often complicated by high rates of mortality and a narrow ototoxic window compared with guinea pig and rat models (40). Prior studies have suggested that Balb/cJ mice have increased susceptibility to cisplatin ototoxicity and reduced mortality compared with CBA/CaJ and C57BL/6J strains (30,41). Owing to its selective high-frequency hearing loss, simplicity, and minimal morbidity, the Balb/cJ strain may be useful for more rapid evaluation and screening of candidate drugs, which can subsequently be tested in a more clinically relevant chronic administration paradigm.

We found that a single IP dose of ISRIB, administered concurrently with cisplatin, significantly reduced cisplatin-induced high-frequency hearing loss by 69%, with near-complete protection of cochlear hair cells. ISRIB, a potent activator of eiF2B and thereby UPR-mediated apoptosis, has been found to effectively modulate the UPR in many disease models, including neurodegeneration (42), memory loss (43), and pulmonary fibrosis (44). We found that ISRIB effectively protects against NIHL (24) and cochlear synaptopathy (25). The current findings suggest that the UPR may be a common mechanism in ototoxicity relating to noise and cisplatin. The efficacy of ISRIB corroborates prior findings: tauroursodeoxycholic acid, a non-specific modulator of the UPR, was effective at mitigating cisplatin ototoxicity *in vivo* (45–46), and salubrinal, which targets eIF2a and, like ISRIB, modulates the PERK/CHOP pathway, protects against cisplatin toxicity *in vitro* (47). Taken together, these findings suggest that targeting the UPR may be a valuable broad strategy to treat acquired hearing loss.

A critical consideration for any potential drug targeting cisplatin ototoxicity is potential interference with cisplatin’s chemoeffectiveness. Indeed, STS, the only currently FDA-approved drug that was shown to be effective in treating cisplatin ototoxicity (5), was also found to be associated with reduced survival in disseminated hepatoblastoma. We found that STS shifts the IC_50_ of cisplatin in a HNSCC cell line by over 10-fold, suggesting that it may indeed interfere with cisplatin’s anti-tumor effects. ISRIB, however, when applied at the same concentration found to reduce CHOP expression (24) and apoptosis (Figure 1) in HEK cells, does not affect the IC_50_ for cisplatin in this cancer cell line. This suggests that specific targeting of the PERK-CHOP pathway of the UPR may be a valuable therapeutic strategy to mitigate cisplatin ototoxicity without interfering with its anti-tumor properties.

### Limitations and Future Directions

This study demonstrates that the UPR is involved in cisplatin ototoxicity; however, several limitations exist. We are unable to resolve which cochlear cell types are involved in this response. RNAscope labelling suggests Chop is upregulated in strial and supporting cells of the inner sulcus and outer sulcus in response to cisplatin treatment, but more work is required to assess cell-type specificity. However, the UPR is a highly conserved mechanism, and we found that HEK cells exhibited similar behavior, including a similar inflection point of the biphasic response. A more significant limitation is that the doses in which the paradoxical response was seen (> 100 μM) are well above those thought to be clinically relevant. However, even if these high cisplatin doses are not clinically relevant, the mechanisms uncovered by these supratherapeutic conditions that lead to hair-cell protection may be relevant when considering drug development for cisplatin ototoxicity.

Despite these *in vitro* limitations, the importance of the UPR in cisplatin ototoxicity is most strongly illustrated by our finding that ISRIB, which inhibits the pro-apoptotic arm of the UPR, can robustly protect against cisplatin-induced high-frequency hearing loss and hair cell death *in vivo*. We used a single-dose acute cisplatin model, which is in contrast with the chronic model of cisplatin administration and ototoxicity that occurs clinically in humans and that has been described in mice (48). Corroboration of this finding in a more clinically relevant chronic administration model of cisplatin ototoxicity would be valuable.

### Conclusion

Cisplatin ototoxicity is a significant challenge with incompletely understood pathophysiology. This study provides evidence for the involvement of the UPR in cisplatin ototoxicity and demonstrates that targeting the pro-apoptotic arm of the UPR with ISRIB mitigates cisplatin-induced hearing loss and hair-cell death in mice without affecting cisplatin chemo-efficacy in a HSNCC cell line. Further investigation of the role of the UPR in cisplatin ototoxicity, and the potential for targeting it for treatment, is warranted.

## Methods

### Sex as a biological variable

Our study examined female mice, because the cisplatin ototoxicity model we used, described below, had less variance in hearing outcomes in female compared with male mice in pilot studies. Therefore, using exclusively female mice in these initial studies reduced the number of animals required. Future studies will examine both female and male mice.

### In vivo mouse model of cisplatin-induced ototoxicity

As a single-dose model for acute cisplatin ototoxicity, 8-week-old female wild-type Balb/cJ mice (Jackson Laboratories, Strain #000651) were administered 5.5 mg/kg cisplatin (Sigma-Aldrich, 1134357 in normal saline) via intraperitoneal (IP) injection. To assess the effect of UPR modulation on cisplatin ototoxicity, we treated animals with ISRIB concurrent with cisplatin administration. Mice were injected intraperitoneally with 10 mg/kg ISRIB (Sigma, SML0843) or vehicle (50% DMSO, 50% PEG-400) immediately before cisplatin administration. Animals were administered 0.5 mL IP normal saline (NS) bolus for hydration concurrent with cisplatin and drug delivery, and subsequently on post injection days 1, 7, 14, and 21.

### Auditory testing

Hearing was tested in mice by measuring auditory brainstem response (ABR) thresholds in response to broadband tone pips at 8, 16, and 32 kHz in the sound field using a standard commercial system (RZ6, Tucker-Davis Technologies) in a soundproof chamber as described (24). All ABR measurements were performed by an investigator blinded to drug treatment.

### Organotypic culture of the neonatal mouse cochlea

Organotypic cultures from wild-type P3-5 C57BL/6J mice (Harlan, 057) were established as described (24). Briefly, the cochlear duct was isolated from P3 mice, opened, and plated on glass coverslips with Cell-Tak (Corning, 354240) with the apical surface of the epithelium facing up. Cultures were used for experiments after 24 hours in culture. Cisplatin was applied to cultures in media in increasing doses of cisplatin from 0-1000 μM for 20 hours.

### UPR gene expression analysis in cell culture

Human embryonic kidney (HEK) cells were grown in DMEM supplemented with 10% FBS, at 37 °C in 5% CO_2_. To measure UPR marker gene expression in response to a broad range of cisplatin doses, HEK cells were treated with cisplatin (0-400 μM) for 40 h, and then total RNA was isolated using TRIzol Reagent (Invitrogen, Carlsbad, CA). 1 μg total RNA was used for first-strand cDNA synthesis using SuperScript IV VILO master mix (Invitrogen, Carlsbad, CA). Real-time PCR was performed by 7900HT FAST Real-time PCR system (Applied Biosystems, Foster City, CA) with a SYBR green I mix. The mRNA primers for UPR markers were as follows: *Chop*: CCACCACACCTGAAAGCAGAA (forward), AGGTGAAAGGCAGGGACTCA (reverse); *S-XBP1*: CTGAGTCCGAATCAGGTGCAG, GTCCATGGGAAGATGTTCTGG; *BiP*: TTCAGCCAATTATCAGCAAACTCT, TTTTCTGATGTATCCTCTTCACCAGT; *DR5*: TGCTGCTTGCTGTGCTACAGGCTGT, TTCTGACAGGTACTGGCCTGCTAG. Relative quantification by the 2^-DDCT^ method was used for analysis as described previously (24). To assess activation of Caspase 3 and 7, the Caspase-Glo^®^ 3/7 assay was performed according to manufacturer’s instructions (Promega) on HEK cells after 40 h treatment with cisplatin. To test if UPR directly causes cisplatin-induced apoptosis, HEK cells treated for 40 h with 0-400 μM cisplatin were additionally treated with 3 doses (at 0, 16 and 24 h) of 0.2 μM ISRIB and the Caspase-Glo^®^ 3/7 assay performed.

### Immunohistochemistry

Evaluation and quantification of hair-cell loss was performed with whole-mount cochlear immunohistochemistry as previously described (24). Briefly, cochlear explant cultures were treated as indicated and fixed in 4% paraformaldehyde. 8-week-old cisplatin-treated mice were euthanized 21 days after cisplatin treatment and temporal bones dissected, isolating the apical, middle, and basal cochlear turns. Specimens were incubated with rabbit anti-myosin7a antibody (a hair-cell-specific marker; 1:200 dilution in PBS; 25-6790, Proteus Biosciences) overnight at 4°C. Cochleae were rinsed three times for 15 min with PBS and then incubated for 2 hours with a goat anti-rabbit secondary antibody conjugated to Alexa Fluor 488 (1:1,000 dilution in PBS; Life Technologies). Whole mounts were rinsed in PBS three times for 15 min and mounted on glass slides in VectaShield antifade mounting medium (Vector Laboratories). The hair-cell region of the organ of Corti was imaged on a Nikon A1R upright line-scanning confocal microscope as previously described. For 8-week-old cochleae, the imaging locations for the apical, middle, and basal turns corresponded approximately to the 8, 16, and 32-kHz regions of the cochlea (25), and missing outer hair cells were counted from anti-Myosin7 immunostain in 200 μm cochlear segments and expressed as a percentage of missing outer hair cells.

### In situ hybridization using RNAScope

To visualize UPR upregulation in the cochlea, we utilized RNAscope to detect CHOP RNA, as a marker of the UPR, and Myosin7a as a hair cell marker in cochlear explants treated with cisplatin. C57BL/6J wild-type mice (Harlan, 057) were dissected as described above and cultured overnight. 100 μM cisplatin was bath applied to cultures for 20 h. Thapsigargin (Sigma), an irreversible inhibitor of SERCA2b which leads to the induction of the UPR, was used as a positive control for induction of the UPR. 1 μM thapsigargin was bath applied to cultures for 2 h, followed by a 1 h fixation in 4% paraformaldehyde.

*In situ* RNA expression analysis was performed using RNAscope Fluorescent Multiplex Reagent Kit (Advanced Cell Diagnostics, Cat. No. 320850) and probes specific to Myosin7a and Chop (RNAscope Probe-Mm-Ddit3-C1, 317661; RNAscope Probe-Mm-Myo7a-C2, 462771-C2) using a protocol adapted from manufacturer’s instructions and previously described protocol for cochlear tissue (49). Cultures were imaged by line-scanning confocal microscopy (Nikon A1R).

### Cell viability in FaDu cells

FaDu cells (gift of Daniel Johnson), a head and neck squamous cell carcinoma line, were cultured in 96-well plates at a density of 10,000 cells per well. Cells were allowed to adhere and grow for 24 h before treatment. Serial dilutions of cisplatin were prepared in cell culture medium at a range of concentrations from 0-200 µM, with 0.2 µM ISRIB, 20 µM z-VAD-FMK, or 3 mM STS. Plates were incubated for 48 h and cell viability assay performed (Promega CellTiter-Glo 2.0). Luminescence was recorded using a SynergyH4 microplate reader (Agilent Technologies, Hayward, CA) and IC_50_ curves analyzed (Prism 9, GraphPad).

### Statistics

For comparison between treatment groups for UPR gene expression, we used one-way ANOVA followed by Tukey multiple comparison tests. For pairwise comparison of ABR thresholds between groups of mice, we used unpaired two-tailed Student’s t-test. Statistical analyses were performed with GraphPad Prism v9.

### Vertebrate studies approval and conduct

All experimental protocols were approved by the UCSF Institutional Animal Care and Use Committee (AN183160). All experiments were performed in accordance with these approved guidelines and regulations; specifically, euthanasia was performed by decapitation for neonatal mice and by CO_2_ euthanasia with cervical dislocation for adult mice. All methods are reported in accordance with ARRIVE guidelines.

## Author Contributions

JL designed and conducted experiments, acquired and analyzed data, and co-wrote the manuscript.

SLR designed and conducted experiments, acquired and analyzed data, and co-wrote the manuscript.

IRM designed and conducted experiments, acquired and analyzed data, and reviewed and revised the manuscript.

EHS designed experiments, analyzed data, reviewed and revised the manuscript, and provided funding.

DKC designed and conducted experiments, acquired and analyzed data, co-wrote the manuscript, and provided funding.

## Acknowledgements

This study was funded by NIDCD R01 DC018583-01 (to DKC and EHS).

